# A machine learning based image classification method to estimate fish sizes from images without a specified reference object

**DOI:** 10.1101/2022.12.27.521854

**Authors:** Catarina NS Silva, Justas Dainys, Sean Simmons, Asta Audzijonyte

## Abstract

Most fish populations around the world are unassessed and their status is unknown. Size based methods could provide fast and transparent assessments, but they require information on fish sizes. Citizen science programs, social media and smart phone applications generate millions of georeferenced fish images globally. Machine learning based fish species and size identification could help turn these images into valuable data for population status assessments. We present a machine learning, image classification based method to identify fish size classes from photos of anglers holding fish. To train the model we group images into ten 5-10 cm size classes, similar to classes used in underwater visual fish surveys. The model was trained using 2602 images from angler citizen science platforms MyCatch and FishSizeProject. Although the number of images was limited, the model achieved an overall accuracy of ~50%. Importantly, the misidentification of size classes was consistent across 20 separate model training rounds, each conducted with an independent, random allocation of images for training and test datasets. Our method suggests that photo based fish size class identification is feasible, and that prediction uncertainty should be incorporated into subsequent analysis, as it is done with fish ageing errors in fisheries stock assessments.

## Method details

### Background

In aquatic ecosystems body size is one of the most important ecological traits, which determines community structure and function [1], population resilience [2], recruitment [3] and species ecological role [4]. Body sizes are also rapidly changing due to warming [5] and harvesting [6]. These changes are likely having a big impact on population dynamics, species interactions, fisheries productivity and food supply for coastal communities [7,8], but remain largely undocumented and unassessed. Fish body size information is also important in size based methods population assessments that provide a way to identify fish population status [9].

Many coastal and freshwater fish species and populations are considered data poor and have no assessment or other information about their status. As a result, they are more likely to be overfished [10]. Yet, the advent of digital technologies means that over the last decade citizen science, angler smart phone applications, social media and other similar platforms have accumulated millions of georeferenced fish images. This data flow is increasing exponentially and has already been used to provide knowledge on invasive species, species redistributions or angler catches [11,12]. However, these data have not been used in more formal population assessments, largely because the relevant information on individual body sizes is lacking. Such lack of population assessments from novel digital data contrasts with another citizen science field of underwater visual surveys, such as Reef Life Survey (https://reeflifesurvey.com/). In these surveys fish body size information is regularly collected by trained volunteer divers, who visually classify fish into 2-10 cm (or larger) size classes. The availability of body size information, even if not precise, means that underwater visual survey datasets have been used in multiple studies of population status assessment [13], including in studies exploring temperature impacts on fish body size [5].

Machine learning methods have already revolutionised research in weather forecasting [14], wildfire disaster response [15], healthcare [16] and transportation [17] and are now providing unprecedented levels of automatic data processing. In fisheries management applications, the uptake has been slower, although automatic fish species identification is now becoming increasing common and is likely to become commonplace. Automatic estimation of fish sizes has also been applied is some cases, but usually requires either a stereo information (video) or clearly defined reference object. Such information is not available in the millions of fish images contributed by anglers and citizen scientists, which are typically characterised by 1) lack a reference object of known length present in the image; 2) the camera is located at an unknown distance from the fish; and 3) the image quality and camera settings differ between images. Yet trained humans usually can estimate an approximate size of fish, which suggest that machine learning approaches may also be successful. In this study we present a new method for fish size identification using machine learning image classification approach and show the feasibility of this method, although the final models need further improvements and training, using considerably large image datasets.

### Data acquisition

To train the model we obtained 3852 multispecies images through a collaborative data sharing agreement between Nature Research Centre and a Canadian angler citizen science platform MyCatch Angler’s Atlas (https://mycatch.ca/). These images have been collected as a part of angler tournaments, where, in addition to a “hero shot” of anglers holding fish, anglers also had to submit a photo of the same fish with a ruler. The tournament rules guaranteed that fish sizes were known precisely. The images with the ruler were only used to confirm the fish sizes and not used in the model training. For quality control we revised all images individually and kept only those images where entire fish body was visible. We also removed images where a ruler was visible in the background, images where fish was held at a very strong angle (e.g. facing the camera), images without a human (only fish was visible), and images with very small (< 10cm) and very large (>80 cm) fish (because such images were rare). This quality control reduced this dataset to 2451 images. In addition, we collected 151 images of known fish sizes through the FishSizeProject smart phone application. The images were collected by Nature Research Centre researchers during regular fish monitoring surveys, which means that fish sizes were also known precisely. The final dataset used for training and testing the model consisted of 2602 images (2451 images from MyCatch Angler’s Atlas and 151 images from FishSizeProject).

Following the Reef Life Survey methodology (https://reeflifesurvey.com/methods/) the final set of images was grouped into 10 sizes classes, using 5 cm size classes for fish from 10 to 40cm, and 10cm classes for fish larger than 40 cm. Like in underwater visual surveys, larger size classes for larger fish account for higher errors associated with sizing large fish [18] (Table 1).

**Table 1:** Number of original images per class and after applying the augmentation technique (vertical and horizontal flip).

| Size class | Original | After augmentation |
| --- | --- | --- |
| class10 | 121 | 363 |
| class15 | 252 | 756 |
| class20 | 190 | 570 |
| class25 | 156 | 468 |
| class30 | 146 | 438 |
| class35 | 165 | 495 |
| class40 | 404 | 1212 |
| class50 | 348 | 1044 |
| class60 | 486 | 1458 |
| class70 | 334 | 1002 |
| Total | 2602 | 7806 |

### Data augmentation

To augment the available image dataset we applied data augmentation techniques. Data augmentation is the process of generating multiple copies of the same images, but with transformations such as rotating, flips, blur, cropping and others. This process has been shown to improve the performance in deep convolutional neural networks [19,20], reduce overfitting [19] and improve model convergence [21].

We used two types of transformations (vertical and horizontal flips), and applied them using the function *A.Compose()* implemented in the Python Albumentations library [22]. The script *data_augmentation_classification.ipynb* available with this publication defines this augmentation pipeline and sets automatic augmentation for multiple images (looping over images, flipping and saving new images).

### Framework

To predict fish sizes in the images we used a simple yet novel machine learning image classification approach. Image classification is different from, for example, machine learning based object detection approach, as it does not require annotating images with bounding boxes, but instead uses the entire image and classifies it into one of the predefined classes, which in our case corresponded to ten fish size classes. Our framework uses the Tensorflow Lite Model Maker library [23,24] and transfer learning, which reduces the amount of training data and time required for model training, compared to a model trained with no prior knowledge. Transfer learning means that we start our image classification model training with a model that is already pre-trained to identify hundreds of different object types. The pre-trained model used here *(efficientnet_lite0* [25]) was trained on the ImageNet dataset [26,27]. As ImageNet includes classes such as “fish”, we hypothesised that this pre-trained model would perform well for fish size classification. The *script fish_size_classification.ipynb* presented with this study uses the Tensorflow Lite Model Maker library to:

1. split the data into training (80%), validation (10%) and testing set (10%)
2. apply the function *image_classifier.create()* to load the pretrained model, load the new image dataset and retrain the model based on the new dataset
3. evaluate model performance using the function *model.evaluate()*
4. generate a confusion matrix for visualising model performance
5. save the trained model.

The parameters used to train the model were ‘*efficientnet_lite0*’ as the pretrained model, batch size of 32 and 10 epochs. The remaining parameters were set to default as defined in the *image_classifier.create()* function. Our initial runs indicated that models with 20 epochs (20 complete passes through all images of the training dataset through the algorithm) initially produced a higher accuracy on the test dataset but had low accuracy on the training dataset. This suggested that the model was overfitted, which can happen when the number of epochs is too large. Depending on the size of the image dataset and the number of classes, we recommend training the model with different number of epochs and evaluating model accuracy using the function *model.evaluate()* in the test and training datasets. Further details on how this can be done are explained in a user friendly code available as an electronic supplement, GitHub and Google Colaboratory environment (see further details in Code Availability section).

### Method validation

The goal of this study was to assess the feasibility of using an image classification framework to identify fish sizes in angler images. Given the complexity of the problem and a relatively large number of size classes (10), it is expected that a well performing model would require some tens of thousands of images and more extensive model training runs. This study only presents a method to develop such a model. Further, to test the replicability and variation in model performance, we repeated the model training procedure 20 times, each time with a random allocation of images into training, validation and testing datasets. (Table 2, Figure 1). The overall accuracy of these models varied between 54% (run 2) and 47% (run 14) and overall loss varied between 1.48 (run 3) and 1.59 (run 14). These replicated model runs aimed to assess whether the classification errors, i.e. allocation of test images into incorrect size classes, are random or partly predictable, such that the model is more likely to assign fishes into neighbouring size classes. To our knowledge such replicated training approach is not typically used in machine learning models, yet it provides a useful and intuitive way to understand whether certain images or a specific allocation of data into training and test datasets can have a large impact on model accuracy. This might be particularly important when computer vision machine learning models are trained on relatively small datasets (up to a few hundred images per class). Our results showed that while there was a substantial variation in the misclassification and prediction accuracy for some size classes (especially 30, 35, 50 and 70, Fig. 1), the model was more likely to misclassify images into neighbouring size classes, as would also be expected with the classification done by humans. Moreover, the replicated training provided a baseline set of misclassification values, which could be incorporated into subsequent statistical analyses of fish size data (e.g. through Bayesian priors).

**Figure 1.**
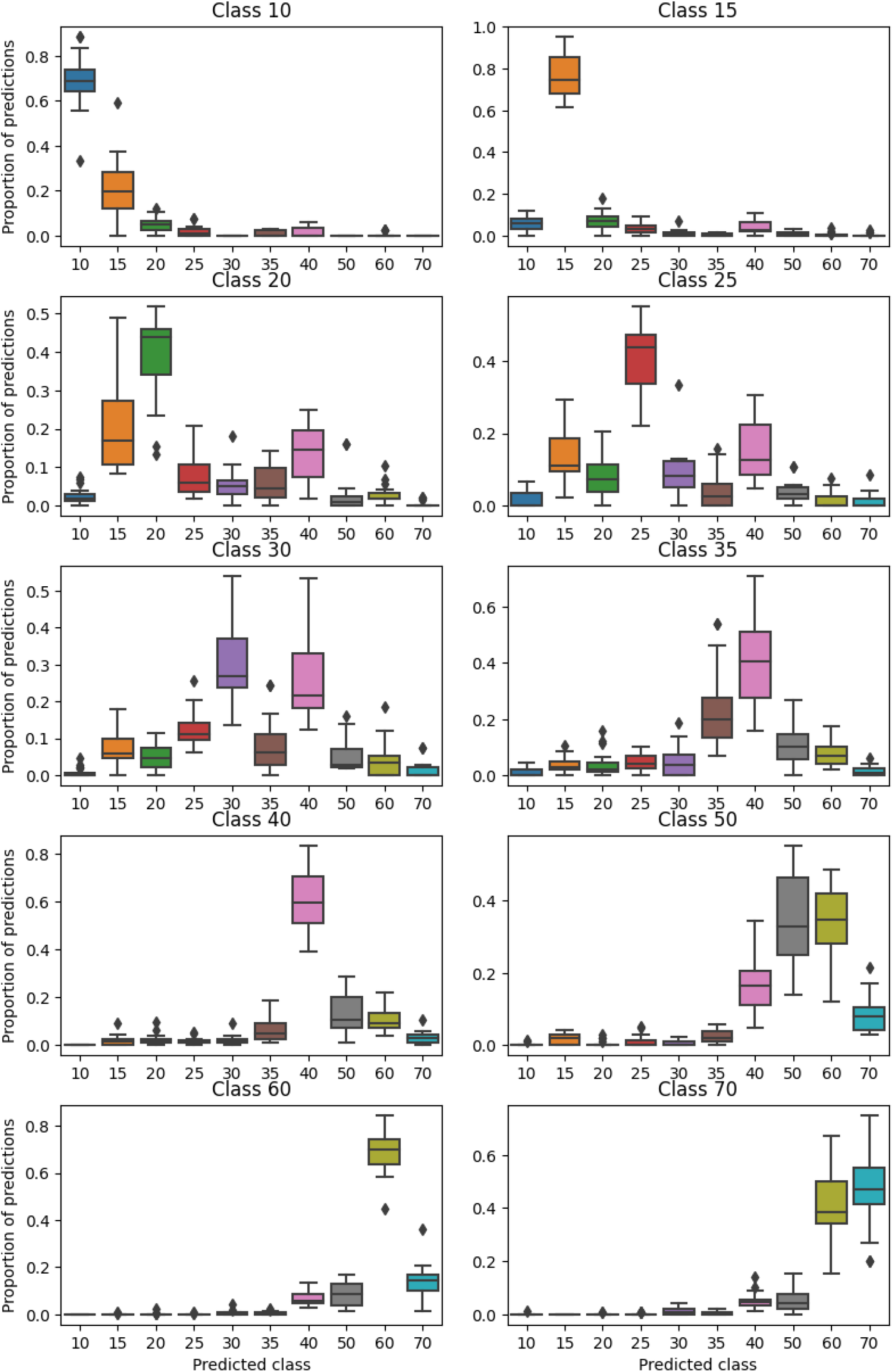
Boxplots of the variation in the proportion of predictions for each class across 20

**Table 2:** Summary of the performance of 20 independent runs for model training (overall accuracy and loss). The model accuracy is a metric that indicates the fraction of predictions where the model was correct (100% indicates a perfectly accurate model). The model loss indicates how bad the model’s prediction was on a single example (the loss is zero if the model’s prediction is perfect).

| Run | Overall Accuracy | Overall Loss |
| --- | --- | --- |
| 1 | 0.52 | 1.55 |
| **2** | **0.54** | **1.50** |
| **3** | **0.54** | **1.48** |
| 4 | 0.52 | 1.56 |
| 5 | 0.53 | 1.54 |
| 6 | 0.52 | 1.55 |
| 7 | 0.51 | 1.53 |
| 8 | 0.50 | 1.57 |
| 9 | 0.51 | 1.55 |
| 10 | 0.53 | 1.57 |
| 11 | 0.50 | 1.54 |
| 12 | 0.54 | 1.55 |
| 13 | 0.53 | 1.50 |
| 14 | 0.47 | 1.59 |
| 15 | 0.50 | 1.53 |
| 16 | 0.52 | 1.52 |
| 17 | 0.53 | 1.52 |
| 18 | 0.49 | 1.58 |
| 19 | 0.52 | 1.53 |
| 20 | 0.51 | 1.53 |

## Visual explanation of the proposed method

Machine learning computer vision models are usually considered “black box” models in that it is often impossible to know which image features are used to make classification decisions. Yet, new methods now allow us to identify the weights that the model assigns to different features in the images. The higher the weight the more important that part of the image is for making classification decisions.

To explore whether the model classified fish sizes based on fishes or other information in the images, we used grad-CAM tool [28] to compute the gradients of the probability in a set of images. Grad-CAM produces a visual explanation using heat-maps that helps understanding how deep learning algorithms make decisions. Figure 2 shows two example images, where parts of the image consistently used in making model predictions are highlighted in red and yellow colours. Regions highlighted in red show that the model considered the fish as the most important area of the image for the classification, which further supported the robustness of the method proposed here.

**Figure 2.**
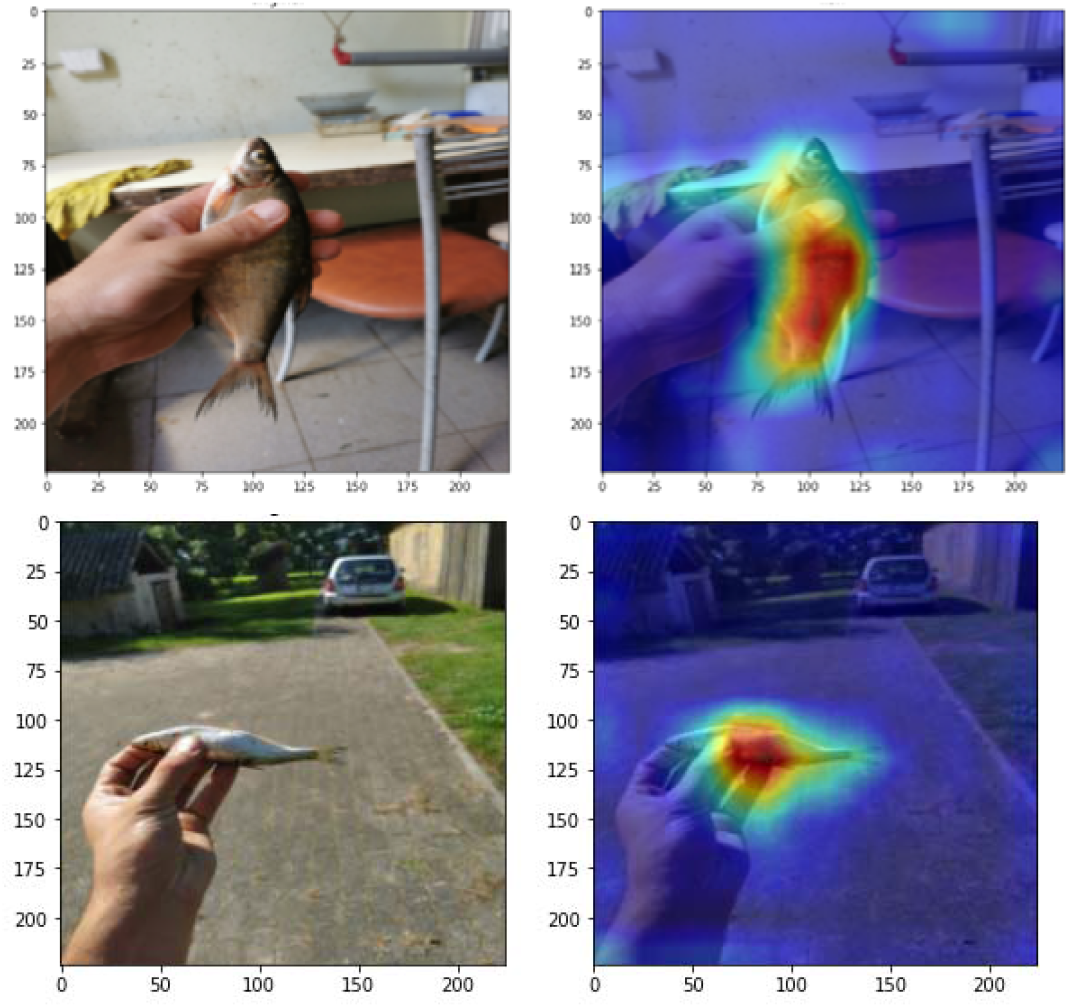
Example heat maps indicating the areas of relevance in the images (in red) that were used as the basis for judgment in image classification. Maps were generated using the Grad-CAM method [28].

## Code availability

The scripts are available as a supplement to this publication and also on the GitHub site https://github.com/fishsizeproject/fish_size_model. The scripts also are uploaded to the Google Colaboratory platform (https://colab.research.google.com/drive/1CIA48l35mdHjRF3jjELWBKHpsDnCs9YE?usp=sharing), which already has many Python libraries installed and provides a limited amount of free computing power for small model runs. To see instructions of how to use these scripts and the Google Colaboratory you can also watch an online course (https://fishsizeproject.github.io/Course-MLforImageProcessing/), where a similar code and framework was applied to a species classification problem.

## Ethics statements

Images in this study were collected through two citizen science platforms MyCatch and scientific research. Images provided together with the example code were anonymised, by removing and blurring all sensitive information.

## CRediT author statement

Catarina NS Silva: Conceptualization, Formal analysis, Methodology, Writing–original draft, Writing–review & editing

Justas Dainys: Data curation, Formal analysis, Writing–review & editing

Sean Simmons: Data Curation, Writing–review & editing

Asta Audzijonyte: Conceptualization, Funding acquisition, Supervision, Methodology, Writing– original draft, Writing–review & editing

## Acknowledgments

This study has received funding from European Regional Development Fund (Project No. 01.2.2-LMT-K-718-02-0006) under grant agreement with the Research Council of Lithuania (LMTLT).

## References

1. De Bie, T.; De Meester, L.; Brendonck, L.; Martens, K.; Goddeeris, B.; Ercken, D.; Hampel, H.; Denys, L.; Vanhecke, L.; Van der Gucht, K.; et al. Body Size and Dispersal Mode as Key Traits Determining Metacommunity Structure of Aquatic Organisms. Ecol. Lett. 2012, 15, 740–747, doi:10.1111/j.1461-0248.2012.01794.x.

2. Savage, V.M.; Gillooly, J.F.; Brown, J.H.; West, G.B.; Charnov, E.L. Effects of Body Size and Temperature on Population Growth. Am. Nat. 2004, 163, 429–441, doi:10.1086/381872.

3. Wootton, H.F.; Audzijonyte, A.; Morrongiello, J. Multigenerational Exposure to Warming and Fishing Causes Recruitment Collapse, but Size Diversity and Periodic Cooling Can Aid Recovery. Proc. Natl. Acad. Sci. U. S. A. 2021, 118, doi:10.1073/pnas.2100300118.

4. Andersen, K.H. Fish Ecology, Evolution, and Exploitation: A New Theoretical Synthesis; Princeton University Press;

5. Audzijonyte, A.; Richards, S.A.; Stuart-Smith, R.D.; Pecl, G.; Edgar, G.J.; Barrett, N.S.; Payne, N.; Blanchard, J.L. Fish Body Sizes Change with Temperature but Not All Species Shrink with Warming. Nat. Ecol. Evol. 2020, 4, 809–814, doi:10.1038/s41559-020-1171-0.

6. Jonathan A. D. Fisher; Frank, K.T.; Leggett, W.C. Breaking Bergmann’s Rule: Truncation of Northwest Atlantic Marine Fish Body Sizes. Ecology 2010, 91, 2499–2505.

7. Oke, K.B.; Cunningham, C.J.; Westley, P.A.H.; Baskett, M.L.; Carlson, S.M.; Clark, J.; Hendry, A.P.; Karatayev, V.A.; Kendall, N.W.; Kibele, J.; et al. Recent Declines in Salmon Body Size Impact Ecosystems and Fisheries. Nat. Commun. 2020, 11, 1–13, doi:10.1038/s41467-020-17726-z.

8. Audzijonyte, A.; Kuparinen, A.; Gorton, R.; Fulton, E.A. Ecological Consequences of Body Size Decline in Harvested Fish Species: Positive Feedback Loops in Trophic Interactions Amplify Human Impact. Biol. Lett. 2013, 9, doi:10.1098/rsbl.2012.1103.

9. Clements, C.F.; Blanchard, J.L.; Nash, K.L.; Hindell, M.A.; Ozgul, A. Body Size Shifts and Early Warning Signals Precede the Historic Collapse of Whale Stocks. Nat. Ecol. Evol. 2017, 1, 1–6, doi:10.1038/s41559-017-0188.

10. Hilborn, R.; Amoroso, R.O.; Anderson, C.M.; Baum, J.K.; Branch, T.A.; Costello, C.; De Moor, C.L.; Faraj, A.; Hively, D.; Jensen, O.P.; et al. Effective Fisheries Management Instrumental in Improving Fish Stock Status. Proc. Natl. Acad. Sci. U. S. A. 2020, 117, 2218–2224, doi:10.1073/pnas.1909726116.

11. Pecl, G.T.; Stuart-Smith, J.; Walsh, P.; Bray, D.J.; Kusetic, M.; Burgess, M.; Frusher, S.D.; Gledhill, D.C.; George, O.; Jackson, G.; et al. Redmap Australia: Challenges and Successes with a Large-Scale Citizen Science-Based Approach to Ecological Monitoring and Community Engagement on Climate Change. Front. Mar. Sci. 2019, 6, 1–11, doi:10.3389/fmars.2019.00349.

12. Sbragaglia, V.; Espasandín, L.; Coco, S.; Felici, A.; Correia, R.A.; Coll, M.; Arlinghaus, R. Recreational Angling and Spearfishing on Social Media: Insights on Harvesting Patterns, Social Engagement and Sentiments Related to the Distributional Range Shift of a Marine Invasive Species. Rev. Fish Biol. Fish. 2022, 32, 687–700, doi:10.1007/s11160-022-09699-7.

13. Edgar, G.J.; Cooper, A.; Baker, S.C.; Barker, W.; Barrett, N.S.; Becerro, M.A.; Bates, A.E.; Brock, D.; Ceccarelli, D.M.; Clausius, E.; et al. Reef Life Survey: Establishing the Ecological Basis for Conservation of Shallow Marine Life. Biol. Conserv. 2020, 252, doi:10.1016/j.biocon.2020.108855.

14. Dewitte, S.; Cornelis, J.P.; Müller, R.; Munteanu, A. Artificial Intelligence Revolutionises Weather Forecast, Climate Monitoring and Decadal Prediction. Remote Sens. 2021, 13, 1–12, doi:10.3390/rs13163209.

15. Jaafari, A.; Zenner, E.K.; Panahi, M.; Shahabi, H. Hybrid Artificial Intelligence Models Based on a Neuro-Fuzzy System and Metaheuristic Optimization Algorithms for Spatial Prediction of Wildfire Probability. Agric. For. Meteorol. 2019, 266–267, 198–207, doi:10.1016/j.agrformet.2018.12.015.

16. Yu, K.H.; Beam, A.L.; Kohane, I.S. Artificial Intelligence in Healthcare. Nat. Biomed. Eng. 2018, 2, 719–731, doi:10.1038/s41551-018-0305-z.

17. Abduljabbar, R.; Dia, H.; Liyanage, S.; Bagloee, S.A. Applications of Artificial Intelligence in Transport: An Overview. Sustain. 2019, 11, doi:10.3390/su11010189.

18. Heenan, A.; Williams, I.D.; Acoba, T.; DesRochers, A.; Kosaki, R.K.; Kanemura, T.; Nadon, M.O.; Brainard, R.E. Long-Term Monitoring of Coral Reef Fish Assemblages in the Western Central Pacific. Sci. Data 2017, 4, 1–12, doi:10.1038/sdata.2017.176.

19. Shorten, C.; Khoshgoftaar, T.M. A Survey on Image Data Augmentation for Deep Learning. J. Big Data 2019, 6, doi:10.1186/s40537-019-0197-0.

20. Mikołajczyk, A.; Grochowski, M. Data Augmentation for Improving Deep Learning in Image Classification Problem. In Proceedings of the 2018 International Interdisciplinary PhD Workshop, IIPhDW 2018; 2018; pp. 117–122.

21. Liu, S.; Papailiopoulos, D.; Achlioptas, D. Bad Global Minima Exist and SGD Can Reach Them. Adv. Neural Inf. Process. Syst. 2020, *2020-Decem*.

22. Buslaev, A.; Iglovikov, V.I.; Khvedchenya, E.; Parinov, A.; Druzhinin, M.; Kalinin, A.A. Albumentations: Fast and Flexible Image Augmentations. Information 2020, 11, 1–20, doi:10.3390/info11020125.

23. Abadi, M.; Barham, P.; Chen, J.; Chen, Z.; Davis, A.; et al,. TensorFlow.Js. In Proceedings of the Proceedings of the 12th USENIX Symposium on Operating Systems Design and Implementation; 2016.

24. Abadi, M.; Agarwal, A.; Barham, P.; Brevdo, E.; Chen, Z.; Citro, C.; Corrado, G.S.; Davis, A.; Dean, J.; Devin, M.; et al. TensorFlow: Large-Scale Machine Learning on Heterogeneous Distributed Systems. arXiv 2016.

25. Tan, M.; Le, Q. V. EfficientNet: Rethinking Model Scaling for Convolutional Neural Networks. 36th Int. Conf. Mach. Learn. ICML 2019 2019, 10691–10700.

26. Russakovsky, O.; Deng, J.; Su, H.; Krause, J.; Satheesh, S.; Ma, S.; Huang, Z.; Karpathy, A.; Khosla, A.; Bernstein, M.; et al. ImageNet Large Scale Visual Recognition Challenge. Int. J. Comput. Vis. 2015, 115, 211–252, doi:10.1007/s11263-015-0816-y.

27. Deng, J.; Dong, W.; Socher, R.; Li, L.-J.; Li, K.; Fei-Fei, L. ImageNet: A Large-Scale Hierarchical Image Database. IEEE Comput. Vis. Pattern Recognit. 2009.

28. Selvaraju, R.R.; Cogswell, M.; Das, A.; Vedantam, R.; Parikh, D.; Batra, D. Grad-CAM: Visual Explanations from Deep Networks via Gradient-Based Localization. Int. J. Comput. Vis. 2020, 128, 336–359, doi:10.1007/s11263-019-01228-7.

